# Microbial Stimulation Reverses the Age-Related Decline in M Cells in Aged Mice

**DOI:** 10.1101/2020.02.17.943514

**Authors:** David S. Donaldson, Jolinda Pollock, Prerna Vohra, Mark P. Stevens, Neil A. Mabbott

## Abstract

Ageing has a profound effect on the immune system, termed immunosenescence, resulting in increased incidence and severity of infections and decreased efficacy of vaccinations. We previously showed that immunosurveillance in the intestine, achieved primarily through antigen sampling M cells in the follicle associated epithelium (FAE) of Peyer’s patches, was compromised during ageing due to a decline in M cell functional maturation. The intestinal microbiota also changes significantly with age, but whether this affects M cell maturation was not known. We show that housing of aged mice on used bedding from young mice, or treatment with bacterial flagellin, were each sufficient to enhance the functional maturation of M cells in Peyer’s patches. An understanding of the mechanisms underlying the influence of the intestinal microbiota on M cells has the potential to lead to new methods to enhance the efficacy of oral vaccination in aged individuals.

## INTRODUCTION

Maintaining the health of an increasingly ageing population is an important challenge for society. The decline in immunity that occurs with ageing, termed immunosenescence, results in decreased vaccine responses and increased incidence and severity of pathogen infections. The intestinal microbiota is required for maturation of the immune system but can also have detrimental effects if it is dysbiotic. Although relatively stable for much of adulthood, ageing induces significant shifts in the composition of the human intestinal microbiota (Claesson et al., 2012). Therefore, restoring immunity in the intestines may have beneficial effects on both intestinal and systemic immune responses.

We have shown that M cell maturation in the intestine is dramatically reduced in aged mice (Kobayashi et al., 2013). M cells are specialised enterocytes normally present in the follicle associated epithelium (FAE) that overlies mucosal lymphoid tissues, such as the small intestinal Peyer’s patches, and equivalents in the nasopharyngeal tract and lung (Date et al., 2017; Kimura et al., 2014; Kimura et al., 2019b; Mabbott et al., 2013). In the intestines, daughter cells derived from Lgr5+ intestinal stem cells at the intestinal crypt base differentiate into M cells upon RANKL stimulation (de Lau et al., 2012) and expression of the transcription factors Spi-B (Kanaya et al., 2012) and Sox8 (Kimura et al., 2019a). M cells transport antigens from the mucosal surface into lymphoid tissues to immune cells that inhabit a unique pocket structure at their basal side (Kolesnikov et al., 2020; Komban et al., 2019). Mice lacking intestinal M cells have delayed development of IgA-secreting plasma cell responses due to impaired germinal centre (GC) formation and T follicular helper (Tfh) cell differentiation (Rios et al., 2016). In the intestine, IgA production is thought to be a critical regulator of the intestinal microbiota and an important mediator against intestinal pathogens. Thus, the delayed IgA response in M cell deficient mice (Rios et al., 2016) highlights the importance of M cells for maintaining a healthy microbiota and in immune responses against pathogens, both of which are reduced with ageing.

Like humans, mice show age-related alterations in their microbiota that are thought to have negative consequences for health. The microbiota is not essential for M cell development, as germ-free mice have similar M cell densities to specific pathogen free (SPF) mice (Kimura et al., 2015). However, other studies have suggested that altering the microbiota may affect M cell development. For example, transferring SPF mice to conventional housing increased the M cell density in Peyer’s patches (Smith et al., 1987). Short-term exposure of rabbit Peyer’s patches to *Streptococcus pneumoniae* was also reported to have a similar effect (Borghesi et al., 1996). Thus, it is possible that reduced M cell maturation in aged mice may be a consequence of changes to the microbiota.

Here, we tested the effect of introducing a faecal microbiota from young mice into aged mice on M cell maturation in small intestinal Peyer’s patches. We found that exposure to a young faecal microbiota restored M cell maturation in aged mice and increased antigen uptake and IgA responses. Furthermore, the M cell density in aged mice could also be restored by stimulation with bacterial flagellin. Cells expressing OLFM4, a stem cell marker, in the intestinal crypts were increased in both conditions suggesting that reduced M cell maturation in aged mice may be a consequence of an age-related decline in intestinal crypt function. By showing that the age-related decline in M cell maturation can be restored, it may be possible to reverse the age-related decline in mucosal vaccine efficacy and the ability to mount protective responses against intestinal pathogens.

## RESULTS

### Passive Microbiota Transfer from Young Donors Enhances M Cell Development in Aged Mice

The gut microbiota changes profoundly during ageing and thus may have an indirect effect on M cell maturation. To explore this further, we facilitated the transfer of the faecal microbiota from young mice into aged mice by housing them for 6 weeks in cages containing used bedding that had previously housed young mice. Groups of control aged mice were housed on clean bedding that had not previously been used to house other mice (**Figure 1A**). After 6 weeks, small intestines were excised from control aged mice, aged mice housed in young bedding, and young donor mice and the Peyer’s patches whole-mount immunostained to detect GP2+ cells, a marker of mature M cells (Hase et al., 2009; Kanaya et al., 2012).

**Figure 1.**
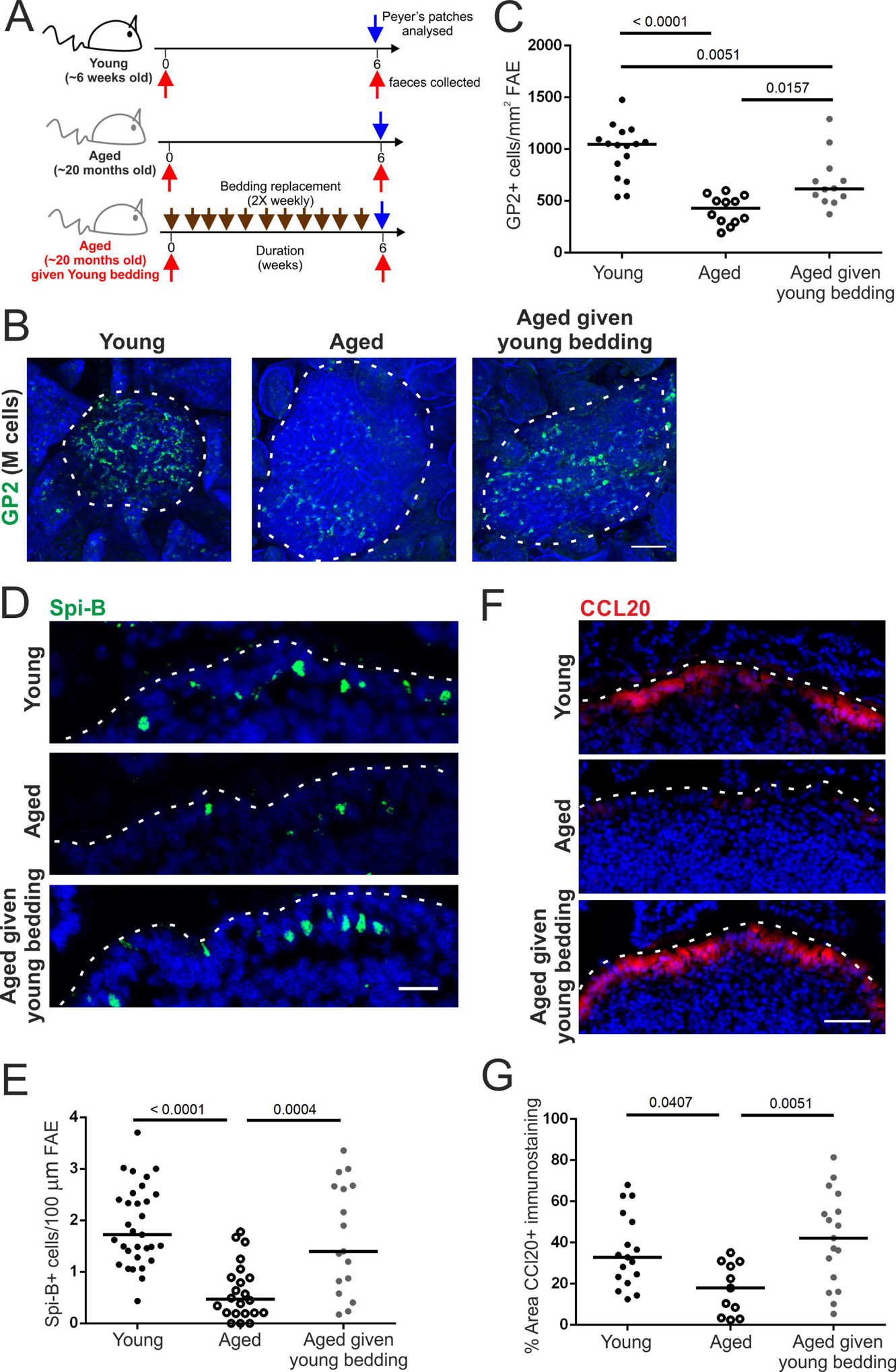
Passive Transfer of a Young Microbiota Enhances M Cell Development in Aged Mice. (A) Cartoon describing the experimental set-up. Aged mice (∼20 months old) were housed for 6 weeks in cages of used bedding that had previously housed young mice. Bedding was replaced twice weekly. Control aged mice were housed in clean cages. (B) Whole-mount immunostaining of GP2+ M cells (green) in Peyer’s patches from young, aged and aged mice given young bedding. Counterstain, F-actin (blue). Scale bar, 100 µm. Broken line, the boundary of the FAE. (C) Quantitation of the number of GP2+ M cells in Peyer’s patches from mice from each group. Each point represents an individual FAE. Horizontal line, median. n=12-16/group from 3-4 mice. Statistical differences determined by one-way ANOVA. (D) IHC detection of Spi-B+ cells (green) in the FAE of Peyer’s patches from mice in each group. Nuclei detected using DAPI (blue). Scale bar, 20 µm. Broken line, the apical surface of the FAE. (E) Quantitation of the number of Spi-B+ cells in the FAE of Peyer’s patches from mice from each group. Each point represents an individual section. Horizontal line, median. n=17-31/group from 3-4 mice. Statistical differences determined by one-way ANOVA. (F) IHC detection of CCL20 (red) in the FAE of Peyer’s patches from mice from each group. Nuclei detected using DAPI (blue). Scale bar, 20 µm. Broken line, the apical surface of the FAE. (G) Quantitation of the % area CCL20+ immunostaining in the FAE of Peyer’s patches from mice from each group. Each point represents an individual FAE section. Horizontal line, median. n=11-17/group from 3-4 mice. Statistical differences determined by one-way ANOVA.

As anticipated, the GP2+ M cell density was significantly reduced in aged mice compared to the young mice. However, in aged mice given young bedding, the GP2+ M cell density was significantly increased, albeit to a significantly lower level found in young mice (**Figures 1B&C**). M cell maturation is dependent on their intrinsic expression of the transcription factor Spi-B (Kanaya et al., 2012). In accordance with the increase in GP2+ M cells, the number of Spi-B+ cells in the FAE was also significantly higher in young mice and aged mice given young bedding (**Figures 1D&E**). This suggests that the decline in M cell maturity in aged mice is in part dependent on age-related changes to the microbiota.

We previously showed that reduced M cell maturity in aged mice was associated with reduced CCL20 expression (Kobayashi et al., 2013), a chemokine produced by the FAE in response to RANKL stimulation. CCL20 is considered to contribute to M cell development by attracting CCR6 expressing lymphocytes to the FAE (Ebisawa et al., 2011). Immunostaining for CCL20 showed a significant reduction in CCL20 in aged mice compared to the young mice (**Figure 1F&G**), consistent with our previous data (Kobayashi et al., 2013). In contrast, aged mice given young bedding had a significantly higher level of CCL20 immunostaining in the FAE compared to control aged mice such that it was indistinguishable from the young mice (**Figure 1F&G**). Therefore, housing aged mice in used bedding from young mice restores CCL20 expression in the FAE.

### Microbiota Transfer from Young Donors Enhances Antigen Uptake and IgA Responses

We next determined if the increased GP2+ M cell density in aged mice given young bedding also increased antigen uptake. Aged mice given young bedding were orally gavaged with 200 nm fluorescent nanobeads, a routinely used model for assessing M cell uptake ability. The number of nanobeads transcytosed into the sub-epithelial dome (SED) region of Peyer’s patches was then determined by microscopy. As anticipated (Kobayashi et al., 2013), significantly more nanobeads were found in the SED of young mice compared to aged mice (**Figures 2A&B**). However, consistent with the increased GP2+ M cell density, aged mice given young bedding had significantly more nanobeads in the SED than control aged mice (**Figures 2A&B**). Indeed, when these data were combined, a significant correlation between M cell density and mean nanobead uptake was observed (**Figure 2C**). The increased nanobead uptake was not associated with changes in mononuclear phagocyte (MNP) populations in the SED. Immunostaining for CD11c revealed similar levels of CD11c+ MNP in the SED of young mice, aged mice and aged mice given young bedding (**Figures 2D&E**). Furthermore, both aged and aged mice given young bedding had reduced numbers of CD11c+ MNP within the FAE (**Figure 2F**). Exposure to young bedding also did not restore the decreased levels of CD68 staining observed in aged mice (**Figures 2D&G**). These data suggest that alterations to the density of MNP populations in the SED were not responsible for the increased bead uptake in aged mice given young bedding.

**Figure 2.**
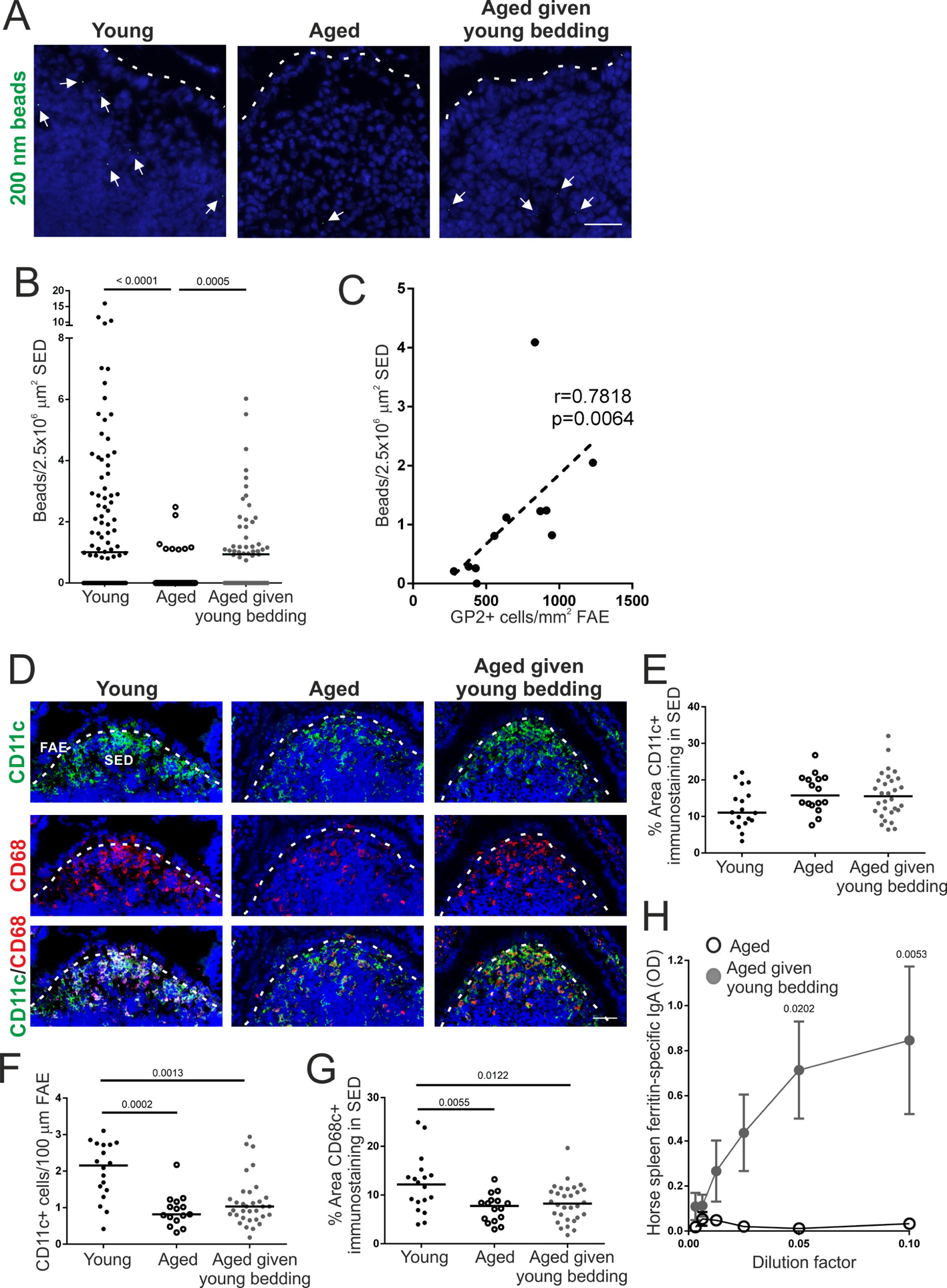
Passive Transfer of a Young Microbiota Enhances Uptake of Particulate Antigen into Aged Peyer’s patches. (A) Histological detection of fluorescent 200 nm nanobeads (arrows) in the SED region of young, aged and aged mice given young bedding. Scale bar, 100 µm. Broken line, the apical surface of the FAE. (B) Quantitation of the number of 200 nm fluorescent nanobeads in the SED of Peyer’s patches from each group. Each point represents an individual section. Horizontal line, median. n=51-94/group from 3-4 mice. Statistical differences determined by Kruskal-Wallis. (C) Corrrelation between GP2+ M cell-density in the FAE and the number of fluorescent 200 nm nanobeads in the SED. Statistical significance determined by Spearman correlation. (D) IHC detection on CD11c+ MNP (green) and CD68+ MNP (red) in the FAE and SED of Peyer’s patches from mice from each group. Nuclei detected using DAPI (blue). Scale bar, 50 µm. Broken line, basal surface of the FAE. (E) The % area of CD11c+ immunostaining in the SED of mice from each group. Each point represents an individual section. Horizontal line, median. n=16-30/group from 3-4 mice. Statistical differences determined by one-way ANOVA. (F) Quantitation of the number of CD11c+ MNP in the FAE of mice from each group. Each point represents an individual section. Horizontal line, median. n=18-33/group from 3-4 mice. Statistical differences determined by Kruskal-Wallis. (G) Comparison of the % area of CD68+ immunostaining in the SED of mice from each group. Each point represents an individual section. Horizontal line, median. n=16-30/group from 3-4 mice. Statistical differences determined by one-way ANOVA. (H) Specific anti-horse spleen ferritin (HSF) IgA levels were determined by ELISA 2 weeks after oral HSF administration in faecal homogenates from aged and aged mice given young bedding. Data presented as mean ± SEM. n=3-4/group. Statistical differences determined by two-way ANOVA.

In the same experiment, aged mice and aged mice given young bedding were administered horse spleen ferritin (HSF) as a model antigen via drinking water as described (Rios et al., 2016), to assess their ability to mount a mucosal antigen-specific IgA immune response. Specific anti-HSF IgA levels were determined by ELISA in faecal homogenates from each group prior to and 2 weeks after HSF administration. When corrected for non-specific IgA levels from samples taken prior to HSF administration, aged mice failed to mount a specific IgA response against this antigen. In contrast, aged mice given young bedding mounted a strong specific IgA response (**Figure 2H**). These data suggest that the increase in M cells in aged mice given young bedding results in increased antigen uptake and consequently an enhanced ability to mount an antigen-specific IgA response to an orally administered antigen.

### M Cell Density and Function Correlates with Increased Abundance of ***Akkermansia municiphila* and Decreased *Turicibacter* Species**

To determine the changes to the intestinal microbiota that occurred following exposure of aged mice to used bedding from young mice, DNA was extracted from faecal pellets from aged mice, aged mice given young bedding and the young donor mice and assessed by 16S rRNA gene metabarcoding. Analysis of molecular variance (AMOVA) testing revealed a significant difference in microbiota structure between young and aged mice (P=0.001). However, no significant difference was observed between aged mice and aged mice given young bedding (P=0.07). Differences in microbiota structure were visualised using non-metric multidimensional scaling (NMDS; **Figure 3A**). Whilst the microbiotas from the young mice formed a tight cluster, both the aged mice and the aged mice given young bedding did not. This suggested greater variation between individuals in the microbiota composition with increased age, confirmed using homogeneity of variance (HOMOVA) testing (P=0.002). Interestingly, no difference in Shannon diversity was observed between the young and aged mice (**Figure 3B**). This indicates that the exposure of aged mice to used bedding from young mice did not result in a large change in microbiota alpha diversity. These data are consistent with data from a similar study of aged C57BL/6 mice exposed to young bedding that also reported that ageing was not associated with a reduction in bacterial diversity (Stebbeg et al., 2019).

**Figure 3.**
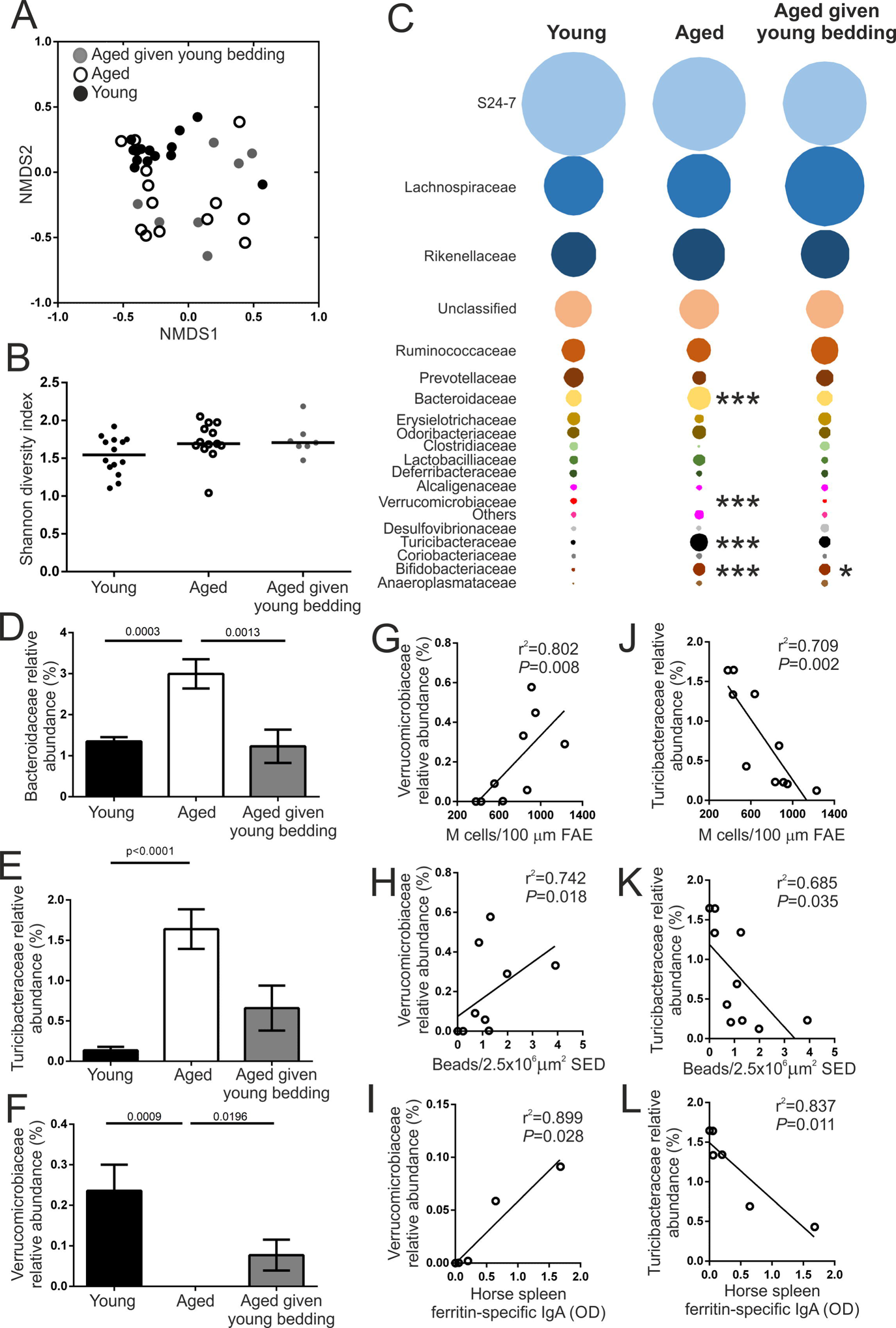
Changes to the Microbiota after housing Aged Mice on Used Bedding from Young Mice. (A) NMDS plot comparing faecal microbiota structure between mice from each age group. n=7-14/group. Faecal microbiotas were significantly different between young mice and each group of aged mice (P=0.002; HOMOVA). (B) Bacterial diversity in faeces from young, aged and aged mice given young bedding was compared using the Shannon index. Horizontal line, median. n=7-14/group. Statistical differences determined by one-way ANOVA (Kruskal-Wallis). (C) Bubble plot showing the relative abundance of distinct bacterial families in the faecal microbiotas of mice from each group. n=7-14/group. Statistical differences for each family determined by one-way ANOVA or Kruskal-Wallis where appropriate. (D-F)The relative abundance of (D) *Bifidobacteriaceae,* (E) *Turicibacteriaceae*, and (F) *Verruocomicrobiaceae* in faeces of young, aged and aged mice given young bedding. Data presented as mean ± SEM. n=7-14/group. Statistical differences determined by Kruskal-Wallis. (G-I) Correlation between the relative abundance of *Verruocomicrobiaceae* in faeces and (G) the density of FAE GP2+ M cells, (H) the number of fluorescent 200 nm nanobeads in the SED, and (I) the level of antigen-specific faecal IgA. Statistical significance determined by Spearman correlation. (J-L) Correlation between the relative abundance of *Turicibacteriaceae* in the faeces and (J) the density of FAE GP2+ M cells, (K) the number of fluorescent 200 nm nanobeads detected in the SED, and (L) the level of antigen-specific faecal IgA. Statistical significance determined by Spearman correlation.

The microbiota was then analysed at family level to look for differences between experimental groups (**Figure 3C**). A significant increase in the relative abundance of *Bifidobacteriaceae* in both groups of aged mice was observed (**Figure 3D**). The relative abundances of both *Bacteriodaceae* and *Turicibacteraceae* were significantly increased with ageing and were reduced in the aged mice given young bedding (**Figures 3D&E**). Conversely, the relative abundance of *Verrucomicrobiaceae* was decreased with ageing and significantly increased in aged mice given young bedding (**Figure 3F**). Of these three bacterial families, only the *Verrucomicrobiaceae* and *Turicibacteraceae* correlated consistently with the functional changes observed in aged mice given young bedding. The relative abundance of *Verrucomicrobiaceae*, represented by the single species *Akkermansia municiphila*, demonstrated a significant positive correlation with M cell density, nanobead uptake and faecal anti-HSF IgA responses (**Figures 3G-I**). Likewise, the relative abundance of *Turicibacteriaceae*, representing a single unclassified *Turicibacter* species, negatively correlated with these parameters (**Figures 3J-L**). These data imply that the decreased abundance of *A. municiphila* and increased abundance of *Turicibacter* species may play a role in the decline in M cell maturation observed in aged mice.

### Systemic Flagellin Treatment Enhances M Cell Development in Aged Mice

We previously showed that the ageing-related decline in M cell maturation was associated with reduced CCL20 expression in the FAE (Kobayashi et al., 2013). The restored functional capacity of M cells in aged mice given young bedding was associated with a restoration of CCL20 expression in the FAE equivalent to that of young mice. Therefore, strategies that boost CCL20 expression in the FAE may directly reverse the decline in M cell maturation in aged mice. Systemic administration of flagellin has been shown to enhance CCL20 expression in the FAE and villous epithelium (Sirard et al., 2009). Furthermore, co-administration of flagellin and nanobeads into the lumen of ligated Peyer’s patches enhanced particle uptake, suggesting a direct modulation of M cell activity (Chabot et al., 2008). Therefore, we hypothesised that systemic flagellin administration may enhance M cell maturation in aged mice.

Aged mice were injected IP with 50 ng of flagellin for 3 d, consistent with similar protocols for enhancing M cell development via systemic RANKL administration (Donaldson et al., 2016; Kanaya et al., 2012; Knoop et al., 2009). Control aged mice were injected with PBS. After 3 d, Peyer’s patches were excised and whole-mount immunostained for GP2. Flagellin administration induced a significant increase in the GP2+ M cell density in aged mice that exceeded the density routinely observed in young C57BL/6J mice in our facility (**Figures 4A&B**). A significant increase in Spi-B+ cells was also observed in the FAE after flagellin treatment (**Figures 4C&D**). Therefore, systemic flagellin administration is sufficient to reverse the decline in GP2+ M cells observed in the FAE of Peyer’s patches from aged mice.

**Figure 4.**
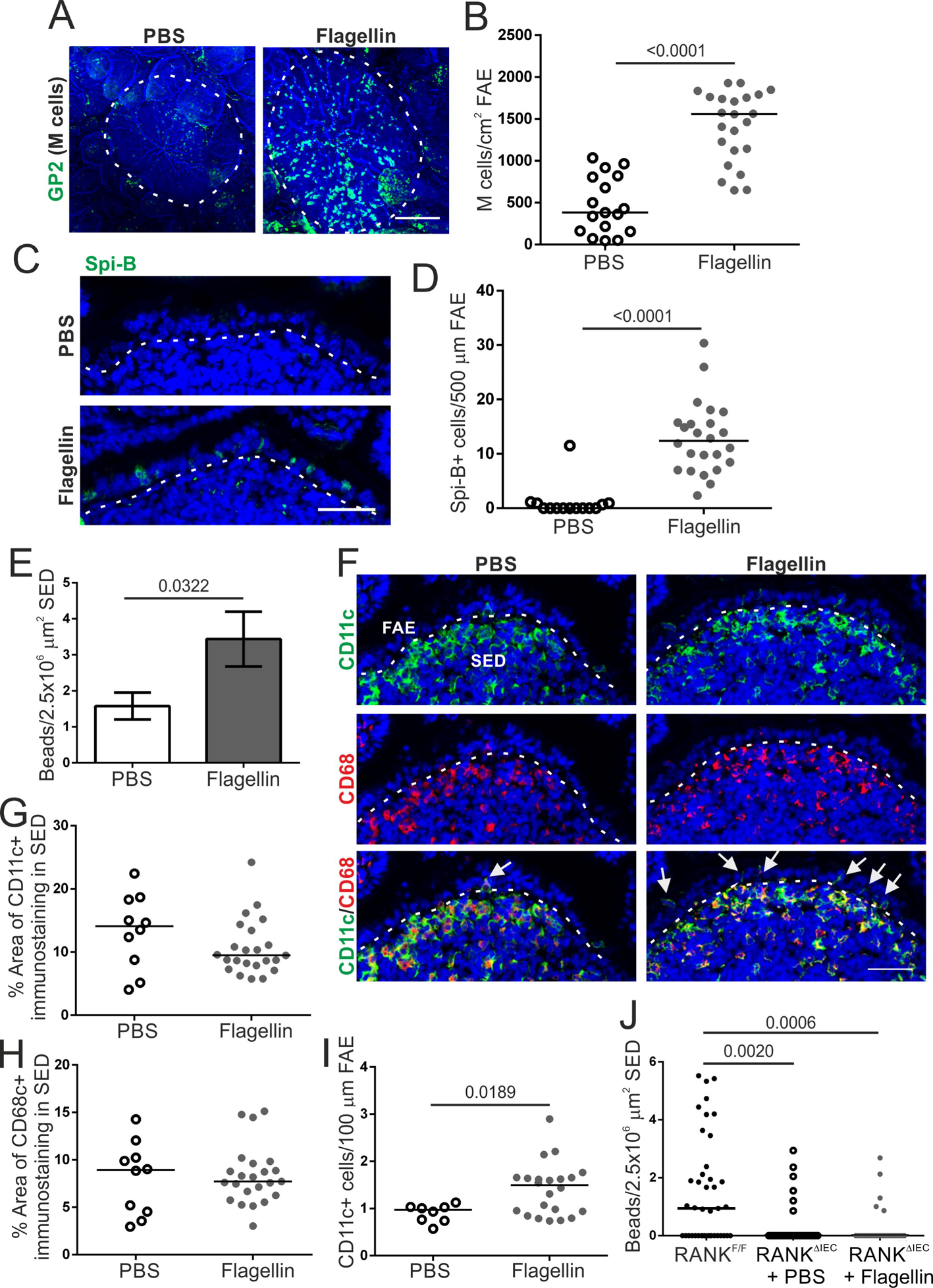
Systemic Flagellin Treatment Enhances M Cell Development in Aged Mice. (A) Whole-mount immunostaining of GP2+ M cells (green) in Peyer’s patches from PBS- and flagellin-treated aged mice. Counterstain, F-actin (blue). Scale bar, 100 µm. Broken line, the boundary of the FAE. (B) Quantitation of the number of GP2+ Mcells in Peyer’s patches from mice from each group. Each point represents an individual FAE. Horizontal line, median. n=17-23/group from 3-4 mice. Statistical difference determined by t-test. (C) IHC detection on Spi-B+ cells (green) in the FAE of Peyer’s patches from mice from each group. Nuclei detected with DAPI (blue). Scale bar, 20 µm. Broken line, the apical surface of the FAE. (D) Quantitation of Spi-B+ cells in the FAE of Peyer’s patches from mice in each group. Each point represents an individual section. Horizontal line, median. n=14-24/group from 3-4 mice. Statistical difference determined by Mann-Whitney. (E) Quantitation of the number of 200 nm fluorescent nanobeads in the SED of PBS- and flagellin-treated aged mice 24 h after orally exposure. Data presented as mean ± SEM. n=53-90/group from 3-4 mice. Statistical difference determined by Mann-Whitney. (F) IHC detection on CD11c+ (green) and CD68+ MNP (red) in the FAE and SED in mice from each group. Nuclei detected with DAPI (blue). Scale bar, 50 µm. Broken line, the basal surface of the FAE. Arrows, CD11c+ cells in FAE. (G-H) The % area of CD11c+ (G) and CD68+ (H) immunostaining in the SED of mice in each group. Each point is from an individual section. Horizontal line, median. n=10-23/group from 3-4 mice. Statistical difference determined Mann-Whitney. (I) Number of CD11c+ cells in the FAE of Peyer’s patches from PBS- and flagellin-treated aged mice. Each point is from an individual section. Horizontal line, median. n=8-23/group from 3-4 mice. Statistical difference determined by t-test. (J) Quantitation of the number of 200 nm fluorescent nanobeads in the SED of PBS or flagellin-treated M cell-deficient RANK^ΔIEC^ mice 24 h after oral exposure. M cell-sufficient RANK^F/F^ mice were used as a positive control. Each point is from an individual section. Horizontal line, median. n=31-41/group from 3 mice. Statistical differences determined by Kruskal-Wallis.

### Systemic Flagellin Treatment Enhances M Cell-Mediated Antigen Uptake in Aged Mice

To confirm that increased M cells in aged mice following flagellin treatment enhanced uptake of luminal antigens, aged mice were treated with flagellin as above and orally gavaged with fluorescent nanobeads on d 3 of flagellin treatment. Nanobead uptake into the SED of Peyer’s patches was compared 24 h later. Consistent with the increase in M cell density, a significant increase in the number of nanobeads in the SED was observed in aged mice treated with flagellin compared to PBS-treated controls (**Figure 4E**). The increased nanobead uptake observed in flagellin-treated aged mice was not due to altered MNP populations within the SED as analysis of CD11c+ and CD68+ immunostaining did not show any difference between PBS and flagellin-treated mice (**Figures 4F-H**). However, the number of CD11c+ cells within the FAE was significantly increased after flagellin treatment, suggesting enhanced interactions between CD11c+ cells and M cells (**Figure 4I**).

To exclude the possibility that the flagellin-mediated increase in particle uptake was due M cell-independent uptake mechanisms such as the increased abundance of CD11c+ cells in the FAE, we took advantage of RANK^Δ^ mice, which have a specific deficiency in RANK expression in the intestinal epithelium and lack M cells (Rios et al., 2016). RANK^Δ^ mice were treated as above with flagellin (or PBS as a control) and orally gavaged with fluorescent nanobeads. M cell-sufficient RANK^FL/FL^ mice were also orally gavaged with nanobeads as a positive control.

Consistent with previous studies (Donaldson et al., 2016; Rios et al., 2016), significantly fewer nanobeads were detected in the SED of Peyer’s patches from flagellin-treated RANK^Δ^ mice compared to RANK mice (**Figures 4J**). Since flagellin treatment did not enhance the uptake of nanobeads in RANK^ΔIEC^ mice, this suggests that the effect of flagellin on antigen uptake into aged Peyer’s patches was due to the flagellin-mediated increase in M cell density.

M cells express a range of receptors on their apical surface that mediate the specific update of distinct pathogenic microorganisms or their toxins (Mabbott et al., 2013). For example, GP2 enables M cells to acquire certain FimH-expressing pathogenic bacteria (Hase et al., 2009). To determine if flagellin treatment also increased M cell-mediated uptake of bacteria, aged mice were treated as above with flagellin (or PBS as a control) and GFP-expressing non-invasive *Escherichia coli* K-12 or invasive *Salmonella* Typhimurium Δ*aroA* (both of which are known to be taken up by M cells via FimH-GP2 interactions) were injected into the lumen of ligated Peyer’s patches. Consistent with the increased uptake of nanobeads, flagellin induced a significant increase in the uptake of *E. coli* K-12 (**Figures 5A&B**) and *S.* Typhimurium Δ*aroA* (**Figures 5C&D**) into the SED of the Peyer’s patches. Dissemination of these bacteria to the mesenteric lymph nodes (MLN) was also determined. As anticipated, the non-invasive *E. coli* K-12 was not detectable in the MLN. However, consistent with the significant increase in *S.* Typhimurium Δ*aroA* uptake in Peyer’s patches of flagellin-treated aged mice, a significantly higher abundance of *S.* Typhimurium Δ*aroA* was recovered from the MLN of the flagellin-treated aged mice (**Figure 5E**). Thus, flagellin treatment enhances the uptake of FimH+ bacteria into Peyer’s patches.

**Figure 5.**
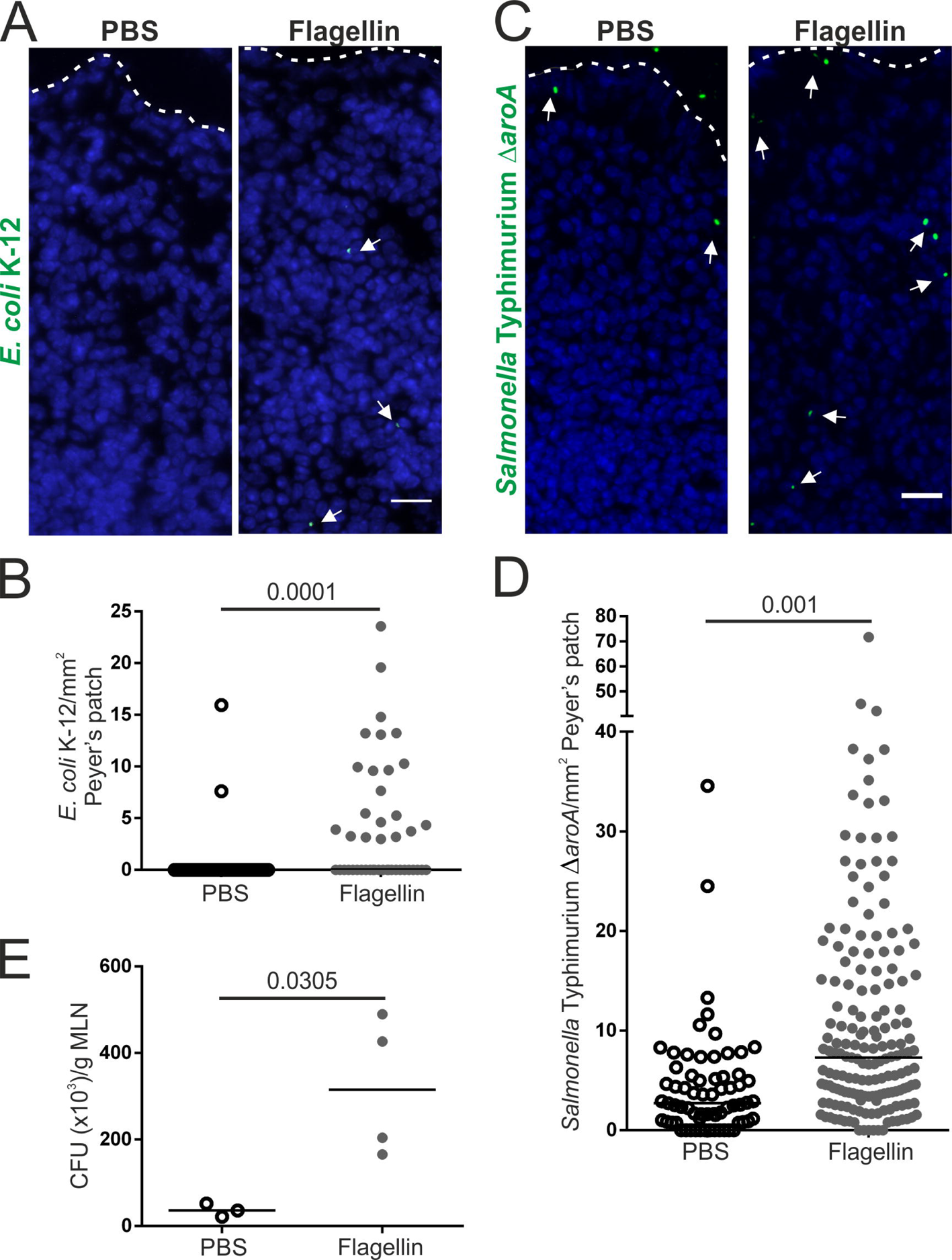
Systemic Flagellin Treatment Enhances Uptake of FimH+ Bacteria into Aged Peyer’s Patches. (A-E) Aged mice (n=3-4) were treated with flagellin (or PBS, control) and GFP-expressing *E. coli* K12 or *Salmonella* Typhimurium injected into ligated loops containing a Peyer’s patch. 1.5 h later the abundance of these bacteria in the SED of Peyer’s patches and MLN was quantified. (A) IHC detection of GFP-expressing *E. coli* K-12 (arrows) in the SED. Nuclei detected using DAPI (blue). Broken line, apical FAE surface. Scale bar, 50µm. (B) Quantitation of the number of GFP-expressing *E. coli* K-12 in the SED of PBS- and flagellin-treated mice. Each point is from an individual section. Horizontal line, median. n=36-45/group from 3-4 mice. Statistical difference determined by Mann-Whitney. (C) IHC detection of GFP-expressing *S.* Typhimurium Δ*aroA* (arrows) in the SED. Nuclei detected using DAPI (blue). Broken line, apical FAE surface. Scale bar, 50 µm. (D) Quantitation of the number of GFP-expressing *S.* Typhimurium Δ*aroA* in the SED of PBS- and flagellin-treated mice. Each point is from an individual section. Horizontal line, median. n=67-171/group from 3-4 mice. Statistical difference determined by Mann-Whitney. (E) Quantitation of *S.* Typhimurium Δ*aroA* colony forming units (CFU)/g in MLN of PBS- and flagellin-treated mice. Points are MLN from individual mice. Horizontal line, median. n=3-4/group. Statistical difference determined by t-test.

### Flagellin Does Not Enhance M Cell Development in Intestinal Organoids *In Vitro*

Although an independent study reported that systemic flagellin-treatment increased CCL20 expression in the FAE (Sirard et al., 2009), our analysis showed no difference in the FAE of aged mice after flagellin treatment (**Figures 6A&B**). We explored this further using *in vitro* enteroids prepared from small intestinal crypts. Whereas RANKL-treatment induced high expression of *Ccl20* mRNA in enteroids (mean 17.4-fold), flagellin-treatment induced only low levels of expression (mean<3-fold) that was not significantly different to untreated enteroids (**Figure 6C**). Furthermore, treatment of enteroids with RANKL and flagellin did not enhance *Ccl20* expression above RANKL-treatment alone (**Figure 6C**). Flagellin was also unable to induce expression of the M cell-related genes *Spib*, *Sox8* or *Gp2* in enteroids, and did not enhance the RANKL-mediated increase in expression of these genes (**Figures 6D-F**). Flagellin stimulates host cells through binding to Toll-like receptor 5 (TLR5). The lack of significant induction of *Ccl20* and M cell-related gene expression in enteroids after flagellin treatment was further supported by the absence of *Tlr5* mRNA expression in deep CAGE sequence data from GP2+ M cells, FAE and RANKL-stimulated enterocytes from the FANTOM consortium (**Figure 6G**; (Forrest, 2014)), and in published mRNA-seq data from isolated GP2+ M cells (**Figure 6H**; (Kimura et al., 2020)). These data suggest that the effect of flagellin on M cell density *in vivo* is not mediated through direct TLR5-mediated stimulation of enterocytes or immature M cells in the FAE.

**Figure 6.**
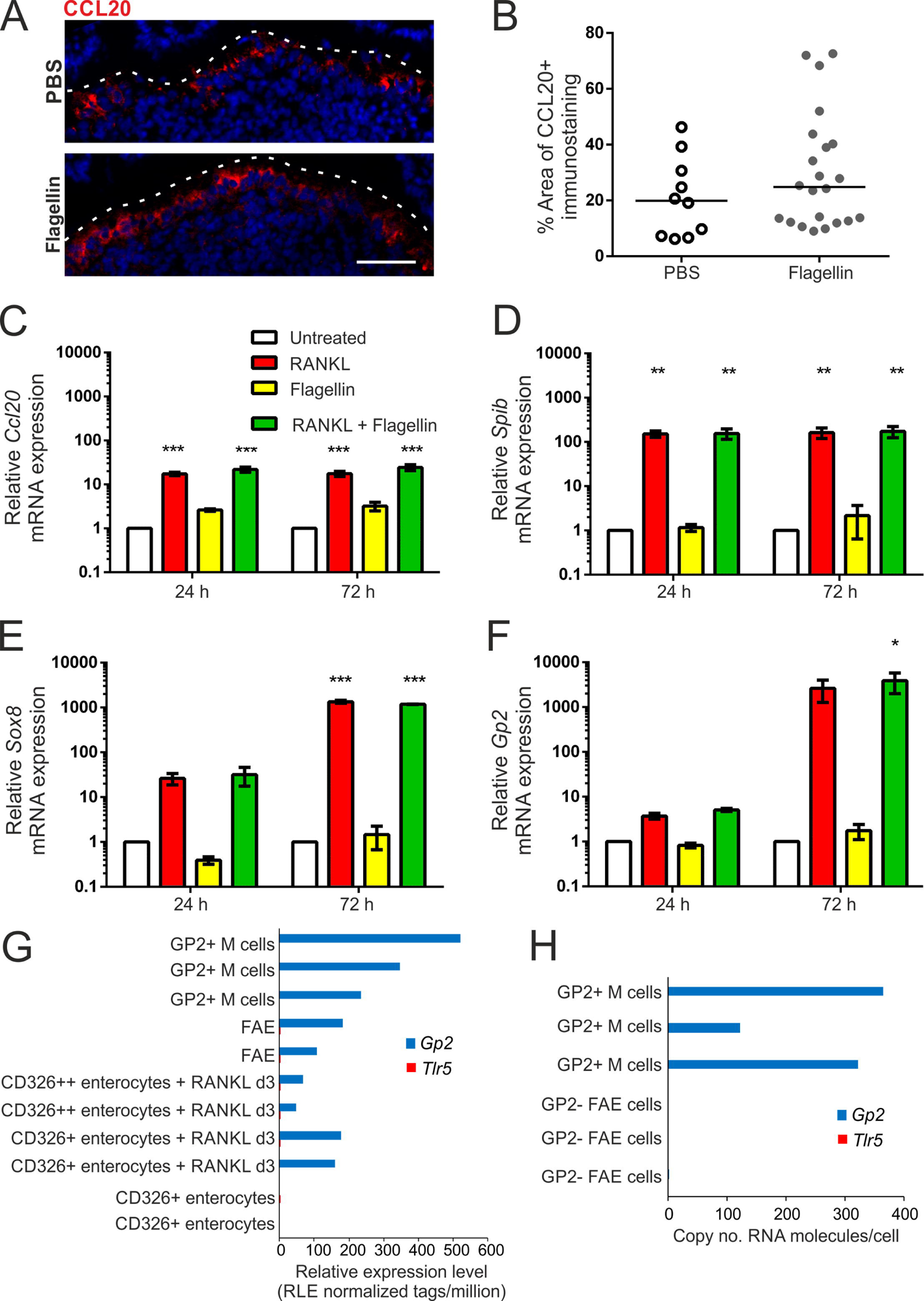
Flagellin Treatment Does Not Enhance M Cell Differentiation in Enteroids. (A) IHC detection of CCL20 (red) in the FAE of Peyer’s patches from PBS- and flagellin-treated aged mice. Nuclei detected with DAPI (blue). Scale bar, 20µm. Broken line shows the apical FAE surface. (B) Quantitation of the % area CCL20+ immunostaining in the FAE of Peyer’s patches from mice in each group of panel A. Each point is from an individual FAE. Horizontal line, median. n=10-22/group from 3-4 mice. Statistical difference determined by Mann-Whitney. (C-F) *In vitro* enteroids prepared from small intestinal crypts were treated with RANKL, flagellin or both. Expression of *Ccl20* (C), *Spib* (D), *Sox8* (E) and *Gp2* (F) was compared by qRT-PCR 24 or 72 h later. Mean expression levels were normalized so that untreated enteroids at 24 h equalled 1.0. Data expressed as mean ± SEM. n=3. Statistical differences determined by two-way ANOVA. (G-H) Comparison of *Tlr5* and *Gp2* mRNA expression (G) in individual cell populations in deep CAGE sequence data from the FANTOM5 project of the FANTOM consortium (Forrest, 2014), and (H) in published mRNA-seq data from isolated GP2+ M cells (GSE108529; (Kimura et al., 2020)).

### Exposure of Aged Mice to a Young Microbiota or Systemic Flagellin-Treatment Restores Small Intestinal Crypts

M cells develop from Lgr5+ intestinal stem cells at the intestinal crypt base (de Lau et al., 2012), and these stem cells are supported by Paneth cells (Sato et al., 2011). Age-related defects in the regenerative ability of intestinal epithelium are thought to arise from impaired Paneth cell function that impairs their ability to support Lgr5+ stem cells (Pentimikko et al., 2019). Interestingly, studies using transgenic reporter mice show that in the small intestinal epithelium, TLR5 expression is restricted to Paneth cells, consistent with data above showing no TLR5 expression in the FAE or M cells. This suggested that the restoration of M cell maturation in aged mice by flagellin may be Paneth cell dependent.

Stimulation of enteroids with flagellin has been shown to increase the expression of olfactomedin-4 (OLFM4) (Price et al., 2018), a marker of intestinal crypt stem cells (van der Flier et al., 2009). Stimulation of our enteroids with flagellin confirmed a significant increase in *Olfm4* mRNA expression (**Figure 7A**). To determine if this occurred *in vivo*, the number of OLFM4+ cells/crypt was determined in sections of small intestine from young, aged and aged mice given young bedding (**Figure 7B**). OLFM4+ cells were observed in both the transamplifying region and the base of the crypt where the Lgr5+ stem cells reside. As anticipated, the total number of OLFM4+ cells/crypt and number of OLFM4+ cells at the base of the crypt of aged control mice was significantly lower than in young mice (**Figures 7C&D**). However, the number of OLFM4+ cells/crypt and at the crypt base was significantly increased in aged mice given young bedding to levels equivalent to young mice. Likewise, a significant increase in OLFM4+ cells/crypt and at the crypt base was observed in flagellin-treated aged mice compared to PBS-treated controls (**Figures 7E-G**). Paneth cell dysfunction in ageing has been linked to increased mTORC activity mice (Pentimikko et al., 2019). Phosphorylated ribosomal protein S6 (pS6) is a downstream effector of mTORC that is increased in aged Paneth cells (**Fig. 7H&I**) (Pentimikko et al., 2019). However, in the crypts of flagellin-treated mice, a significant reduction in the area that stained positive for pS6 was observed (**Fig. 7H&I**). Together, these data suggest that the increased M cell maturation in aged mice induced by both young bedding and flagellin was due to restored Paneth cell function and their effects on stem cells within the intestinal crypts.

**Figure 7.**
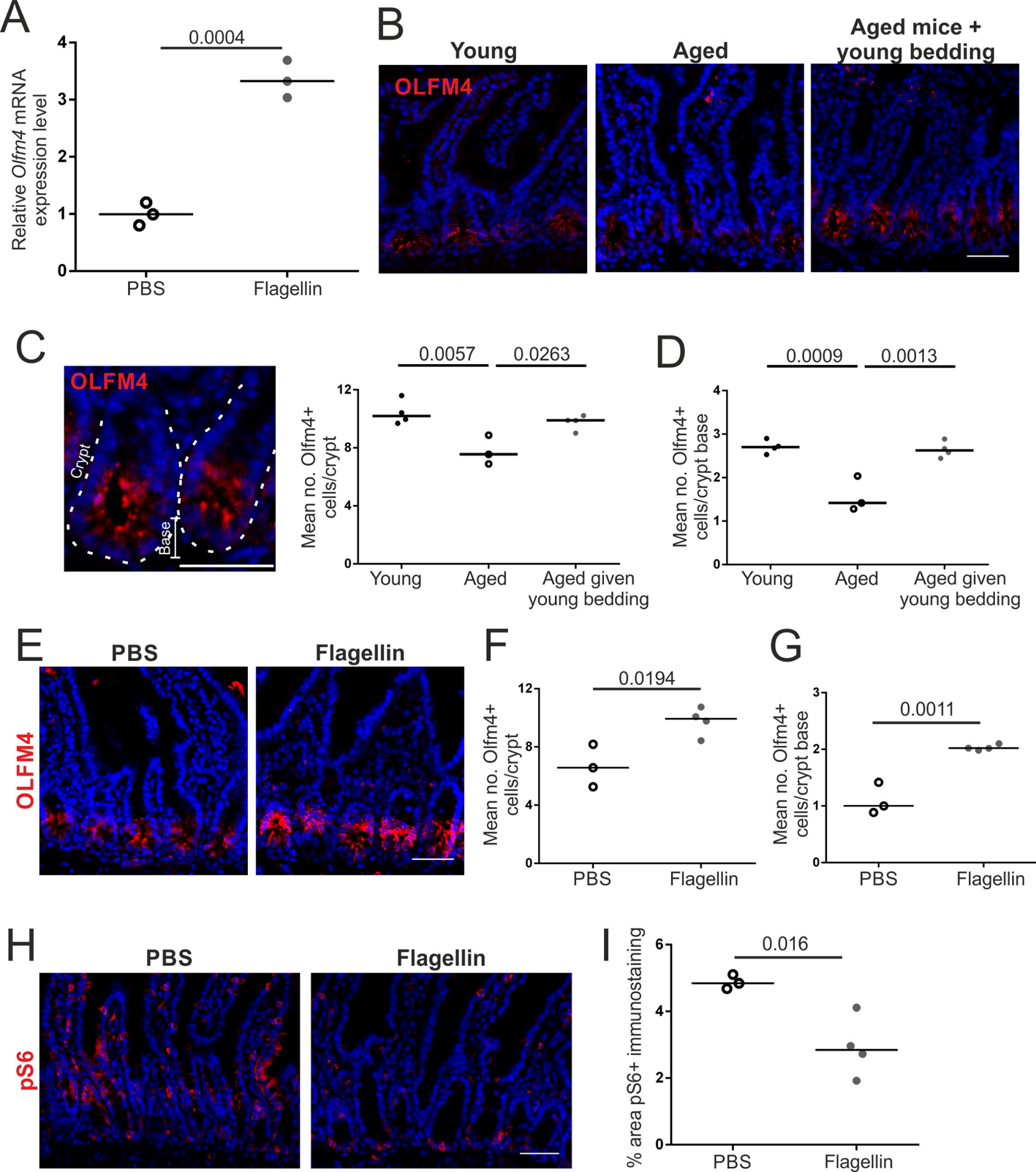
Exposure of Aged Mice to a Young Microbiota or Flagellin Restores Small Intestinal Crypts. (A) Olfm4 mRNA expression was compared by qRT-PCR 24 h after flagellin stimulation of *in vitro* enteroids prepared from small intestinal crypts. Horizontal line, median. n=3/group. Statistical difference determined by t-test. (B) IHC detection of OLFM4 (red) in small intestinal crypts of young, aged and aged mice given young bedding. Nuclei detected with DAPI (blue). Scale bar, 50 µm. (C-D) Quantitation of the number of OLFM4+ cells in small intestinal crypts (C) and the crypt base (D) from mice in each group in panel B. Each point is the mean of an individual mouse. n=3-4/group. Horizontal line, median. Statistical differences determined by one-way ANOVA. (E) IHC detection of OLFM4 in small intestinal crypts from PBS- and flagellin-treated aged mice. Nuclei detected with DAPI (blue). Scale bar, 50 µm. (F-G) Quantitation of the number of OLFM4+ cells in small intestinal crypts (F) and the crypt base (G) from mice in each group in panel E. Each point is the mean of an individual mouse. n=3-4/group. Horizontal line, median. Statistical differences determined by by t-test. (G)IHC detection of pS6 (red) in small intestinal crypts from PBS- and flagellin-treated aged mice. Nuclei detected with DAPI (blue). Scale bar, 50 µm. (H) Quantitation of the % area pS6+ immunostaining in small intestinal crypts of mice in each group in panel H. Each point is the mean of an individual mouse. n=3-4/group. Horizontal line, median. Statistical differences determined by t-test.

## DISCUSSION

The decline in immunity with age significantly affects the health and well-being of the elderly. Here, we show that the age-related decline in M cell maturation can be restored by flagellin stimulation or exposure to a young faecal microbiota. The restoration of M cell maturation in aged mice resulted in enhanced antigen uptake and intestinal IgA responses against a model antigen. M cells develop from stem cells located in the intestinal crypts (de Lau et al., 2012). Interestingly, both transfer of a young microbiota and flagellin stimulation increased the numbers of OFLM4+ cells in the crypt, suggesting that improving crypt function may be important in re-establishing intestinal immunity in ageing.

Mice lacking M cells have impaired faecal IgA responses (Rios et al., 2016), highlighting the important role of antigen sampling by M cells in the development of the immune response against the microbiota. The age-related decline in M cell maturation is likely to affect the development of IgA responses against the microbiota, demonstrated by aged mice failing to mount a specific IgA response against HSF. Therefore, the age-related decline in M cell maturation may contribute to the age-related changes in the microbiota. The aged microbiota contributes to age-associated gut inflammation (Fransen et al., 2017). Thus, reversing the decline in M cell maturation may have beneficially alter the aged microbiota composition.

IgA responses depend on the development of GC responses in the Peyer’s patch, and GC B and Tfh cells are also reduced in M cell deficient mice (Rios et al., 2016). Consistent with the decline in M cell maturation (Kobayashi et al., 2013), Peyer’s patch GC B and Tfh cells are also reduced in aged mice (Stebbeg et al., 2019). The decline in Peyer’s patch GC B and Tfh cells in aged mice could be reversed by transfer of a young microbiota (Stebbeg et al., 2019), consistent with the results of this study. It is therefore likely that the increased GC B and Tfh cells observed in aged mice after young microbiota transfer is dependent on increased M cell maturation.

M cell maturation results in the ability to transcytose antigens. Increased antigen trafficking into the Peyer’s patch would stimulate naïve lymphocytes. This could be enhanced by activated antigen-specific B cells that can sample antigens directly from M cells and migrate to the GC (Komban et al., 2019). Additional mechanisms beyond antigen transcytosis could also contribute to increased IgA production and GC reactions in aged mice. M cells also sample nanoparticles that capture microbiota-derived immune-stimulatory macromolecules such as peptidoglycan (Powell et al., 2015). Interestingly, this drives PD-L1 expression in the Peyer’s patch, which may signal to Tfh cells through PD-1, an absence of which alters the GC response and results in poor IgA selection (Kawamoto et al., 2012). Additionally, other bacterial ligands may be important in stimulating Tfh development and IgA responses. For example, germ-free mice containing T cells that lack TLR signalling have reduced Tfh cell development in response to a TLR2 agonist and fail to mount specific intestinal IgA responses (Kubinak et al., 2015). Therefore, bacterial ligands transported into Peyer’s patches of aged mice alongside antigens could effectively act as adjuvants that boost the GC reaction and specific IgA development.

Our analysis of the changes to the faecal microbiota that occurred in aged mice given young bedding revealed a positive correlation between M cells, bead uptake and specific IgA response with the abundance of *A. municiphila*, and a negative correlation between the same parameters and an unclassified *Turicibacter* species. Further studies are required to determine if these directly affect the mucosal immune system or intestinal crypts, or are merely representative of broader changes to the microbiota associated with the restoration of immunity. Interestingly, aged mice fed resistant starch, as a means to increase short chain fatty acid (SCFA) production, also show reduced *Turicibacter* and increased *A. municiphila* abundance (Tachon et al., 2013) suggesting a reciprocal link between abundance of these and SCFA levels, known dampeners of intestinal inflammation.

The abundance of *A. municiphila* is also decreased in aged humans (Collado et al., 2007), and reduced *A. municiphila* is associated with the development of inflammation in germ-free mice reconstituted with an aged microbiota (Fransen et al., 2017). *A. municiphila* is a mucin degrading bacterium (Derrien et al., 2008) and colonic mucus thickness has been shown to be reduced with age (Elderman et al., 2017). An association between reduced small intestinal mucin-producing goblet cells in aged mice and the abundance of *A. municiphila* has also been reported (Sovran et al., 2019). It also helps in maintain epithelial integrity *in vitro* (Reunanen et al., 2015). *A. municiphila* also upregulates genes involved in metabolism in enteroids potentially through SCFA production (Lukovac et al., 2014). A recent study also noted that nicotinamide produced by *A. municiphila* was protective in a model of amyotrophic lateral sclerosis (Blacher et al., 2019). Nicotinamide is a subunit of nicotinamide adenine dinucleotide (NAD^+^). Mouse enteroids treated with a NAD+ precursor, nicotinamide riboside, restores aged enteroid growth *in* vitro (Igarashi et al., 2019). Interestingly, nicotinamide riboside also restores the reduction in crypt OLFM4+ cells *in vivo* (Igarashi et al., 2019) as seen in our aged mice given young bedding, suggesting a link between *A. municiphila* produced nicotinamide and M cell restoration in aged mice.

Little is known of the unclassified *Turicibacter* species, however *Turicibacteriaceae* are decreased in mice that lack TNFα (Jones-Hall et al., 2015) and increased in humans with rheumatoid arthritis (Chen et al., 2016). Increased TNFα production may underlie macrophage dysfunction and microbiota dysbiosis in the ageing intestine (Thevaranjan et al., 2017). Thus, the decreased *Turicibacter* observed in this study may be indicative of decreased ageing-associated intestinal inflammation. Whether reduced age-related inflammation promotes increased M cell maturation or is a consequence of increased M cell maturation is not known.

Although previous studies have suggested that systemic flagellin treatment stimulates CCL20 expression throughout the intestinal epithelium (Sirard et al., 2009), CCL20 was not significantly increased in the FAE in our study. Consistent with this, the ability of flagellin to induce CCL20 in enteroid cultures was modest in comparison to that induced by RANKL. In the intestinal epithelium TLR5 is predominantly expressed by Paneth cells (Price et al., 2018) which help maintain intestinal crypt stem cells. We have shown that macrophage depletion results in an altered Paneth cell phenotype that impairs M cell development (Sehgal et al., 2018). Paneth cell function is also defective in the ageing intestine (Pentimikko et al., 2019). The increase in OLFM4+ cells observed in the intestinal crypts of aged mice treated with flagellin is likely due to direct stimulation of TLR5-expressing Paneth cells. However, other mechanisms, such as stimulation of haematopoietic cells (Kinnebrew et al., 2010), may also synergise with this to promote the large increase in M cell maturation observed after flagellin treatment.

How flagellin stimulation increases OLFM4 expression by stem cells in aged mice is unknown, but may improve wnt signalling, as exogenous wnt signalling restores stem cell function in aged enteroids (Nalpareddy et al., 2017). Similar effects are observed after notum inhibition, a wnt inhibitor upregulated in aged Paneth cells (Pentimikko et al., 2019). Interestingly, flagellin stimulation increases expression of genes associated with mTORC1 signalling in enteroids (Price et al., 2018). Ageing Paneth cells also have higher expression of genes associated with mTORC1 signalling, and inhibition of this pathway restores intestinal crypt cell function (Pentimikko et al., 2019). Following flagellin stimulation, pS6, a downstream effector of mTORC, was reduced in the intestinal crypts suggesting that mTORC activity was reduced by flagellin stimulation. Although seemingly contradictory, this suggests that flagellin activation of mTORC may initiate autoregulatory pathways that inhibit mTORC1 activity (Yang et al., 2019). This may underlie the effect of flagellin on Paneth cells and the subsequent effects on M cell maturation and intestinal immunity.

In conclusion, our data suggest that the age-related decline in intestinal immunity can be reversed by boosting M cell numbers through manipulation of the microbiota or flagellin stimulation. Restoring the M-cell to IgA axis in the elderly could offset the harmful effects associated with the age-related changes to the microbiota and thus improve health. Additionally, this could be used to enhance responses to oral vaccination or improve the outcome of intestinal pathogen infections to which aged individuals are more susceptible. The observation that this may rely on improving intestinal crypt stem cell function means that treatments aimed at restoring the regenerative capacity of the aged intestine by modulating intestinal stem cells may have the added benefit of improving intestinal immunity.

## ACKNOWLEDGEMENTS

We thank Barry Bradford, Dave Davies, and Bob Fleming for excellent technical support. This work was supported by funding from the Biotechnology and Biological Sciences Research Council (grant numbers BB/M024288/1; BBS/E/D/20002174 & BBS/E/D/30002276) and Medical Research Council (grant number MR/S000763/1).

## AUTHOR CONTRIBUTIONS

N.A.M. conceived the study and obtained funding; D.S.D. and N.A.M. designed the study; D.S.D, J.P. and P.V. performed the experiments; J.P. helped design and perform the microbiota analyses; P.V. and M.P.S. provided GFP-expressing bacteria, helped design and perform the bacteria uptake experiments; D.S.D. and N.A.M. wrote the manuscript; all authors contributed to the writing of the final version of the manuscript.

## DECLARATIONS OF INTEREST

The authors declare no competing interests.

## METHODS

### Mice

Female C57BL/6J mice were purchased from Charles River (Margate, UK). RANK^Δ^ and RANK mice (Rios et al., 2016) were bred at the University of Edinburgh. Mice were maintained in-house under specific pathogen-free conditions to the ages required. Young mice were used at 6-8 weeks old, aged mice were used at approx. 20-26 months old. All the experiments described in this study were first approved by The Roslin Institute’s Ethical Review Committee, and were conducted under the authority of a UK Home Office project licence in full compliance with the Animals (Scientific Procedures) Act 1986.

### Passive microbiota transfer

To facilitate the passive transfer of the faecal microbiota from young mice to aged mice, we housed aged mice for a 6 week period in cages containing used bedding that had previously been used to house young mice. To provide the donor bedding, young mice were removed from their caging and placed into a fresh cage with clean bedding. The aged mice were then housed in the empty used cage that previously been used to house the young mice. This was repeated twice weekly (every 3-4 d) for a 6 week period. Two cages of donor young mice were used and the aged mice were alternated between them. Groups of aged mice were housed on clean bedding that had not previously been used to house other mice as a control.

### Bacterial 16S rRNA Gene Metabarcoding

Faecal samples were collected from aged mice before and at 4 and 6 wk after passive microbiota transfer and from young donor mice. DNA was extracted and prepared for 16S rRNA gene sequencing, targeting the V3 hypervariable region, as described previously (Pollock et al., 2018). One library pool was constructed using equimolar concentrations of DNA from each of the included samples (n=39), as calculated using a fluorometric assay (Qubit dsDNA HS Assay kit, Thermo Fisher Scientific, Paisley, UK). Additionally, a mock bacterial community (20 Strain Even Mix Genomic Material ATCC®MSA-1002, ATCC, USA), a reagent-only control sample and two sham samples (empty tube controls exposed only to air in the room at the time of sampling) were included in the pool to assess sequencing error rate and background DNA contamination. Using the mock bacterial community data suggested the sequencing error rate was ∼0.01%.

The library pool was quantified using the Quant-iT™ PicoGreen® double-stranded DNA Assay Kit (Thermo Fisher Scientific) to ensure adequate DNA yield for sequencing using the Illumina MiSeq (Illumina, Cambridge, UK) using V2 chemistry and generating 250 bp paired-end reads (Edinburgh Genomics, Edinburgh, UK). The primer sequences were first removed from the forward and reverse reads using cutadapt (Martin, 2011). The sequence files generated with the primers removed are publicly available through the European Nucleotide Archive (ENA) under the project accession number PRJEB36358.

Mothur (version 1.40.5) (Schloss et al., 2009) was then used to generate contiguous sequences, and to carry out sequence quality control and analysis as described by the software developers (URL: https://www.mothur.org/wiki/MiSeq_SOP; accessed February 2019). Unique sequences were binned using a database-independent approach. A mean of 140,836 sequences were obtained per sample after quality control, with one sample being removed from the analysis due to a low number of sequences being retained. Files were subsampled to the lowest number of sequences obtained (n = 4101) for analysis. The Shannon Index was calculated for each sample to assess alpha diversity. To assess beta diversity, a distance matrix was constructed using Yue and Clayton theta similarity coefficients (Yue and Clayton, 2005). To visualise community similarities between groups, Non-Metric Multidimensional Scaling (NMDS) plots were compiled. The statistical significance of differences in clustering by treatments was assessed by analysis of molecular variance (AMOVA) (Excoffier et al., 1992). The statistical significance of variation between populations was tested using homogeneity of molecular variance (HOMOVA) (Stewart Jr. and Excoffier, 1996).

### Systemic bacterial flagellin treatment

Mice were given 10 µg Ultrapure flagellin from *S.* Typhimurium (Invivogen, Toulouse, France) in sterile PBS by intra-peritoneal injection daily for 3 d.

### *In vivo* Uptake of Fluorescent Nanobeads

Mice were given a single oral gavage of 2 × 10^11^ of Fluoresbrite Yellow Green labelled 200 nm microbeads (Polysciences, Hirschberg an der Bergstrasse, Germany) in 200 μl PBS. Mice were culled 24 h later and Peyer’s patches were snap-frozen in liquid nitrogen. Serial frozen sections (6 μm in thickness) were cut on a cryostat and counterstained with DAPI (4′,6-Diamidine-2′-phenylindole; Thermo Fisher Scientific). The number of beads in the SED from 3-4 sections of two Peyer’s Patches per mouse (*n*=3–4 mice/group; total 9–33 SED/mouse studied) were counted. Images of Peyer’s patches SED regions were acquired using Nikon Eclipse E400 fluorescent microscope using Micro Manager (http://www.micro-manager.org). Tissue auto-fluorescence was subtracted from displayed images using ImageJ.

### Bacterial Strains and Culture Conditions

An *aroA* deletion mutant of *Salmonella* enterica serovar Typhimurium 4/74 (ST4/74 Δ*aroA*; (Buckley et al., 2010)) and *E. coli* K-12 strain DH4α were routinely cultured at 37°C in Luria-Bertani (LB) broth and on LB agar. ST4/74 Δ*aroA* was cultured with 50 µg ml^-1^ kanamycin. To aid visualization of infected mouse cells *in vivo*, both strains were electroporated with the plasmid pFPV25.1, which carries *gfpmut3A* under the control of the *rpsM* promoter resulting in the constitutive synthesis of GFP (Valdivia and Falkow, 1996). Following electroporation, strains were routinely cultured at 37°C in media supplemented with 100 µg ml^-1^ ampicillin to maintain pFPV25.1. The plasmid is known to be stable in *Salmonella* in bovine ileal loops *in vivo* over 12 h (Vohra et al., 2019).

### Bacterial uptake into Ligated Peyer’s Patches

For inoculation of murine ligated ileal loops, overnight cultures of GFP-expressing ST4/74 Δ*aroA* and *E. coli* K-12 were diluted to obtain approximately 10^9^ colony forming units (CFU)/ml. Viable counts were confirmed retrospectively by plating of 10-fold serial dilutions of the cultures on LB agar containing 100 µg/ml ampicillin.

Mice were anesthetised and a gut loop prepared centred on an individual Peyer’s patch. Each loop was inoculated with 100 µl of culture (∼ 10^8^ CFU). Mice were culled 1.5 h later and Peyer’s patches from the loops snap-frozen in liquid nitrogen. Serial frozen sections (6 μm thickness) were cut on a cryostat and counterstained with DAPI. The number of GFP-expressing E. coli K-12 in the SED were counted directly in 6 sections of Peyer’s patch per mouse (n=3–4 mice/group; total 9–16 SED/mouse studied). GFP-expressing ST4/74 Δ*aroA* was visualised by immunostaining with rabbit polyclonal anti-GFP and Alexa594 labelled anti-rabbit IgG (both Thermo Fisher Scientific). GFP-expressing ST4/74 Δ then counted in 12 sections of Peyer’s patch per mouse (n=3–4 mice/group; total 9– 56 SED/mouse studied). Images of Peyer’s patches SED regions were acquired using Nikon Eclipse E400 fluorescent microscope using Micro Manager (http://www.micro-manager.org). Tissue auto-fluorescence was subtracted from displayed images using ImageJ software (https://imagej.nih.gov/ij/).

### Bacteriological analysis of tissues

Mesenteric lymph nodes (MLNs) from infected mice were snap-frozen in liquid nitrogen. Tissues were thawed and then gently washed in PBS to remove non-adherent bacteria and weighed. A 10% homogenate was prepared in PBS using a Tissue Lyser II (Qiagen, Manchester, UK) and stainless steel beads. Ten-fold serial dilutions were plated both on LB agar and MacConkey agar containing 100 µg/ml ampicillin with or without 50 µg/ml kanamycin to differentiate between ST4/74 Δ and E. coli K-12.

### Oral Immunization with Horse Spleen Ferritin

Antigen-specific faecal IgA responses to an orally administered antigen were assessed in aged mice as previously described (Rios et al., 2016). Four weeks after the passive microbiota transfer was initiated, horse spleen ferritin (1mg/ml; Sigma, Gillingham, UK) was orally administered to aged mice via drinking water on days 0–2 and 7–9. Faecal samples were collected 2 wk later and a 10% homogenate (w/v) prepared in PBS. Horse spleen ferritin-specific IgA levels in the supernatant of the faecal homogenates was determined by enzyme-linked immunosorbent assay in plates coated with horse spleen ferritin followed by detection with horseradish peroxidase-conjugated goat anti-mouse IgA (Southern Biotech, Birmingham, AL, USA) using optEIA (BD Biosciences, Oxford, UK) as the substrate. O.D. values were corrected for background using matched fecal samples collected immediately prior to the commencement of oral immunisation.

### IHC Analysis

To detect M cells by whole-mount immunostaining Peyer’s patches were first fixed using BD Cytofix/Cytoperm (BD Biosciences), and then immunostained with rat anti-mouse GP2 mAb (MBL International, Woburn, MA). Peyer’s patches were then stained with Alexa Fluor 488-conjugated anti-rat IgG Ab and Alexa Fluor 647-conjugated phalloidin to detect F-actin (both Thermo Fisher Scientific).

Peyer’s patches and small intestines were also snap-frozen at the temperature of liquid nitrogen, and 6μm serial frozen sections cut using a cryostat. To detect MNP, sections were immunostained with hamster anti-mouse CD11c mAb (clone N418, Thermo Fisher Scientific) and rat anti-mouse CD68 mAb (clone FA-11, BioLegend, London, UK). To detect Spi-B, paraformaldehyde-fixed frozen sections were treated with citrate buffer (pH 7.0, 121°C, 5 min) before immunostaining with sheep anti-mouse Spi-B polyclonal Ab (R&D Systems, Abingdon, UK). CCL20, OLFM4 and pS6 were detected in paraformaldehyde-fixed frozen sections using goat anti-mouse CCL20 polyclonal Ab (R&D Systems), rabbit anti-mouse OLFM4 mAb (clone D6Y5A0) or rabbit anti-phospho-S6 ribosomal protein mAb (clone D68F8) (both Cell Signalling Technology, London, UK). Sections were subsequently immunostained with species-specific secondary antibodies coupled to Alexa Fluor 488 (green) or Alexa Fluor 594 (red) dyes (Thermo Fisher Scientific). Cell nuclei were detected using DAPI. Sections were mounted in fluorescent mounting medium (DAKO, Stockport, UK) prior to imaging on a Zeiss LSM710 confocal microscope (Zeiss, Cambourne, UK).

### Image Analysis

Digital microscopy images were analysed using ImageJ software as described previously (Inman et al., 2005). Background intensity thresholds were first applied using an ImageJ macro which measures pixel intensity across all immunostained and non-stained areas of the images. The obtained pixel intensity threshold value was then applied in all subsequent analyses. Next, the number of pixels of each color (black, red, green, yellow etc.) were automatically counted and presented as a proportion of the total number of pixels in each area under analysis. To analyse immunostaining in FAE and SED regions, 3-8 images were routinely analysed/mouse. For wholemounts, GP2+ cells were counted in 4-7 FAE/mouse. Cell counting in sections of FAE (Spi-B+ cells, CD11c+ cells) was performed on 1-15 images/mouse. The number of OLFM4+ cells was counted in 28-74 crypts/mouse.

### *In Vitro* Enteroid Cultivation

Intestinal crypts were dissociated from mouse small intestine using Gentle Cell Dissociation Reagent (Stemcell Technologies, Cambridge, UK). The crypts were then re-suspended in Intesticult medium (Stemcell Technologies) at 4×10^3^ crypts/ml and mixed 1:1 with Matrigel (Corning, Flintshire, UK). Next, 50 μl Matrigel plugs were plated in pre-warmed 24-well plates and allowed to settle, before addition of 600 μl of prewarmed Intesticult medium and subsequently cultured at 37°C in a 5% CO_2_ atmosphere. Fresh medium was replaced every 2 d of cultivation and the enteroids passaged after 7 days of culture. Where indicated, enteroids prepared from the 1st passage were treated with either RANKL (50ng/ml, Biolegend), ultrapure flagellin from *S.* Typhimurium (100ng/ml; Invivogen) or in combination. For each experimental condition, enteroids were cultivated in triplicate and repeated using enteroids from three independent animals.

### Real-time Quantitative PCR (RT–qPCR) Analysis of mRNA Expression

For mRNA extraction, enteroids were incubated in Cell Recovery Solution (Corning) for 1 h at 4 °C. Total RNA was then isolated using RNeasy Mini Kit (Qiagen) followed by removal of genomic DNA and cDNA synthesis using SuperScript IV VILO Master Mix with ezDNase Enzyme (Thermo Fisher Scientific) both as per manufacturer’s instructions. PCR was performed using the Platinum-SYBR Green qPCR SuperMix-UDG kit (Thermo Fisher Scientific) and the Stratagene Mx3000P real-time qPCR system (Stratagene, CA, USA). The following qPCR primers were used: *Ccl20* (5’-CGACTGTTGCCTCTCGTACA-3’ and 5’-AGCCCTTTTCACCCAGTTCT-3’); *Gapdh* (5’-GGGTGTGAACCACGAGAAAT-3’ and 5’-CCTTCCACAATGCCAAAGTT-3’); *Gp2* (5’-GATACTGCACAGACCCCTCCA-3’ and 5’-GCAGTTCCGGTCATTGAGGTA-3’); *Olfm4 (5’-* TGGCCCTTGGAAGCTGTAGT-3’ and 5’-ACCTCCTTGGCCATAGCGAA-3’); *Sox8* (5’-TCCGTTGCTCTCCGGTTT-3’ and 5’-GCCCATTCTCTCCTTTGTCCT-3’); *Spib* (5’-AGCGCATGACGTATCAGAAGC-3’ and 5’-GGAATCCTATACACGGCACAGG-3’).

### Statistical Analyses

Details of all group/sample sizes and experimental repeats are provided in the figure legends. Statistical analyses were performed in Prism 6 (Graphpad Software, San Diego, CA). Details of tests used are provided in the figure legends. In instances where there was evidence of non-normality (identified by the D’Agostino & Pearson omnibus, Shapiro-Wilk or Kolmogorov–Smirnov normality test), data were analysed using appropriate non-parametric tests. Values of P < 0.05 were accepted as significant.

